# BET inhibition prevents aberrant RUNX1 and ERG transcription in STAG2 mutant leukaemia cells

**DOI:** 10.1101/762781

**Authors:** Jisha Antony, Gregory Gimenez, Terry Taylor, Umaima Khatoon, Robert Day, Ian M. Morison, Julia A. Horsfield

## Abstract

Mutations in the subunits of the cohesin complex, particularly in the STAG2 subunit, have been identified in a range of myeloid malignancies, but it is unclear how these mutations progress leukaemia. Here, we created isogenic K562 erythromyeloid leukaemia cells with and without the known leukemic STAG2 null mutation, R614*. STAG2 null cells acquired stem cell and extracellular matrix gene expression signatures that accompanied an adherent phenotype. Chromatin accessibility was dramatically altered in STAG2 null K562 cells, consistent with gene expression changes. Enhanced chromatin accessibility was observed at genes encoding hematopoietic transcription factors, ERG and RUNX1. Upon phorbol 12-myristate 13-acetate (PMA)-induced megakaryocytic differentiation, STAG2-null cells showed precocious spike in RUNX1 transcription from its P2 promoter. A similar precocious spike was observed in transcription of ERG. Interestingly, spikes in RUNX1-P2 and ERG only occurred as immediate early response to differentiation induction. Treatment of STAG2 null cells with enhancer-blocking BET inhibitor, JQ1, dampened precocious RUNX1 P2 expression and led to a complete loss of RUNX1 P1 and ERG transcription during PMA stimulation in both parental and STAG2 null K562 cells. These results suggest that precocious RUNX1 and ERG expression in STAG2 null cells is enhancer-driven. Furthermore, JQ1 treatment reduced stem cell-associated KIT expression in STAG2 null cells. We conclude that STAG2 depletion in leukemic cells amplifies an enhancer-driven transcriptional response to differentiation signals, and this characteristic is dampened by BET inhibition. The results have relevance to the development of therapeutic strategies for myeloid leukaemia

---

Cohesin is a multiprotein complex that is essential for cell division but also has key roles in genome organisation that underpin its gene regulatory function. Recurrent mutations of genes encoding cohesin subunits occur in myeloid malignancies at 10-12% (Kon et al., 2013), and the frequency of cohesin mutation in Down-Syndrome associated megakaryoblastic leukaemia (DS-AML) is even higher (∼50%) (Yoshida et al., 2013). Cohesin insufficiency reinforces stem cell programmes and impairs differentiation in haematopoietic stem cells (HSC) (Mazumdar et al., 2015; Mullenders et al., 2015; Viny et al., 2015). The STAG2 subunit of cohesin is the most frequently mutated in myeloid malignancies (Kon et al., 2013). In contrast to other cohesin subunits, complete loss of STAG2 is tolerated due to partial compensation by STAG1. STAG2 and STAG1 have redundant roles in cell division (Benedetti et al., 2017; van der Lelij et al., 2017). However, cohesin-STAG1 and cohesin-STAG2 have non-redundant roles in facilitating 3D genome organization to delineate tissue specific gene expression (Kojic et al., 2018).

Cohesin depletion was previously shown to alter chromatin accessibility and transcription of the *RUNX1* and *ERG* genes (Mazumdar et al., 2015), which encode transcription factors that regulate haematopoietic differentiation. Here we used CRISPR-Cas9 to edit K562 erythroleukaemia cells to contain a patient-specific STAG2 R614* mutation (Mullenders et al., 2015) and found that *RUNX1* and *ERG* are precociously transcribed in response to phorbol 12-myristate 13-acetate (PMA)-induced megakaryocytic differentiation.

We characterised two K562 edited lines with homozygous STAG2 R614* mutation (*STAG2-null*^*A*^, *STAG2-null*^*B*^) (Figure 1A and Supplementary Figures 1-2; Supplementary Material). Both *STAG2-null* lines showed complete loss of STAG2 (Figure 1B). *STAG2-null* K562 cells exhibited occasional adherent characteristics (Figure 1C) and slower cell cycle progression (Supplementary Figure 3). Array CGH showed that both *STAG2-null* lines had varying minor gains and losses of genetic material relative to the parental line (Supplementary Figure 4). Nevertheless, both *STAG2-null* transcriptomes clustered together and were distinct from the parental line (Supplementary Figure 5). Consistent with potential compensation by STAG1, both *STAG2-null* lines showed 1.6-fold upregulation in *STAG1* (Supplementary Figure 6). Several transcription factors, kinases, chemokines, cytokines and lineage markers that were lowly expressed in parental cells were significantly upregulated in one or both *STAG2-null* clones (Supplementary Figure 7). Gene set enrichment analyses revealed a loss of the typical K562 associated chronic myelogenous transcription profile (Supplementary Figure 8). *STAG2-null* cells upregulated extracellular matrix genes reflecting their adherent phenotype, and gained a stem cell-like expression signature (Figure 1D and Supplementary Figure 8). These results show that STAG2 depletion leads to profound morphological and transcriptional changes.

**Figure 1.**
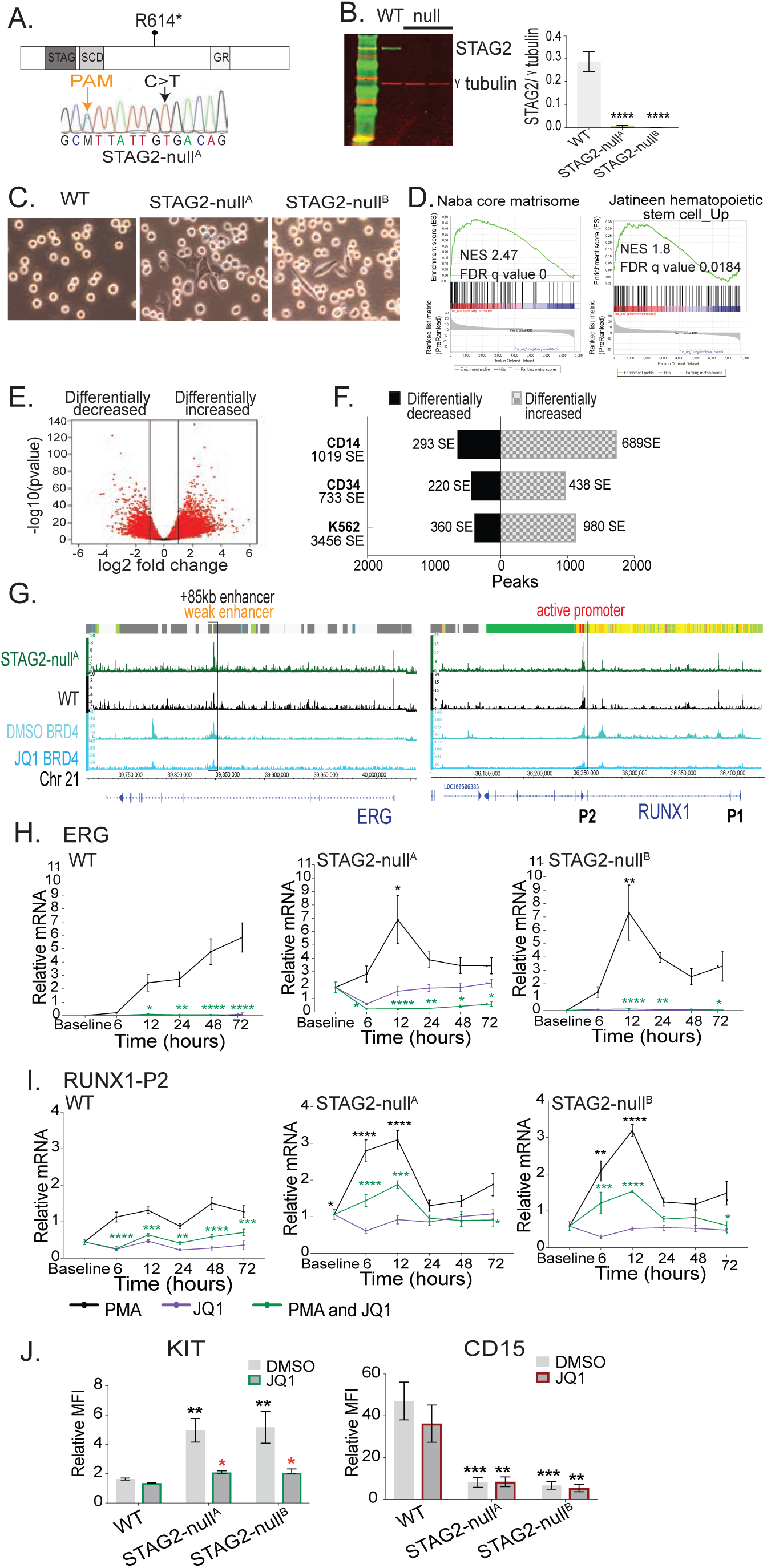
STAG2 mutation alters chromatin accessibility and response to cell-signalling. (A) Schematic of STAG2 protein showing the position of STAG2 R614* (C>T) mutation. Shown also is the Sanger sequencing plot for edited K562 cells containing homozygous STAG2 R614* mutation (*STAG2-null*^*A*^). A silent mutation was introduced at PAM site in STAG2 mutants. (B) Immunoblot analyses of STAG2 protein levels in parental (WT) and STAG2 mutant cells. Bar graphs show STAG2 protein normalized to γ tubulin from 3 biological replicates. Significance was determined by unpaired t-test. (C) Images of parental (WT) and *STAG2-null* K562 cells in culture (D) Gene set enrichment analyses showing upregulation of extracellular matrix (Naba core matrisome) and haematopoietic stem cell genes in *STAG2-null*^*A*^. Shown are the normalised enrichment score (NES) and FDR-q value (E) Volcano plot of differential chromatin accessibility in *STAG2-null*^*A*^ compared to WT K562 cells. Significant peaks at adjusted p-value ≤ 0.05 are shown in red (52,452 sites showed differential accessibility, 29,432 differentially increased and 23,020 differentially decreased). Lines indicate log2 fold change cut off-2. (F) Enrichment of differentially increased and decreased accessible sites identified in *STAG2-null*^*A*^ at super enhancers (SEs; defined in K562, CD34+ cord blood cells and CD14+ monocytes). (G) Integrative genome browser view of normalized ATAC-sequencing signals from *STAG2-null*^*A*^ and WT cells at *ERG* and *RUNX1*. Significant (p ≤ 0.05) accessible sites at *RUNX1-P2* promoter and *ERG* +85 enhancer are boxed. ChromHMM data for K562 (derived from ENCODE) is shown at the top of each plot, and additional tracks are BRD4 binding in K562 following treatment with DMSO or 6 hours of JQ1 (Liu et al. 2017). (H) *ERG* and (I) *RUNX1-P2* expression levels examined over a time course treatment with PMA, JQ1 or a combination of PMA and JQ1. Graphs depict average relative mRNA levels from 3 biological replicates normalized to 2 reference genes. Black asterisks denote significant difference between WT and *STAG2-null* lines following PMA only treatment. Green asterisks denote significant difference between PMA only and combination of PMA and JQ1 treatment within each cell type. Significance was determined by two-way Anova. (J) Relative mean fluorescence intensity (MFI) of KIT and CD15 following treatment with control DMSO or JQ1 for 24 hours. Relative MFI for each cell type and condition was determined as a ratio of MFI in stained/unstained. Graphs represent the average of 3 biological replicates. Significance was determined by two-way Anova. Black asterisks denote a significant difference between parental (WT) and *STAG2-null* cells for the same condition. Red asterisks denote a significant difference between DMSO and JQ1 treatment within each cell type. (*p<0.05, **p<0.01, ***p<0.001, ****p<0.0001).

ATAC-sequencing showed that chromatin accessibility was differentially altered at ∼50,000 sites in *STAG2-null*^*A*^ cells (Figure 1E). Motif analyses of differentially accessible sites identified strong enrichment for the enhancer-regulating bZIP or AP-1 factors (FRA1, FRA2, JUN-AP1) at sites of increased accessibility, and for CTCF and CTCFL (BORIS) at sites of decreased accessibility (Supplementary Figure 9). In *STAG2-null* cells we observed increased chromatin accessibility at super-enhancers (SEs) defined for K562, CD34+ primary cord blood cells and CD14+ monocytes (Figure 1F). 45% genes near SEs with differential accessibility also displayed altered transcript levels in *STAG2-null*^*A*^ cells. SE-proximal genes included those encoding cell lineage marker or transcription factors (Supplementary Figure 10).

The *RUNX1* and *ERG* loci contain SEs in CD34+ cells. SEs in proximity to *RUNX1* and *ERG* gained accessibility in *STAG2-null*^*A*^ cells (Supplementary Figure 11). Many of the increased accessible sites were bound by a variety of AP-1 factors at *RUNX1*, and primarily by JUND at *ERG* (Supplementary Figure 11). Closer visualization revealed that the prominent ATAC sites in K562 are at the stem cell-associated *ERG* +85 kb enhancer and at *RUNX1*-P2 promoter, and both these sites showed increased accessibility in *STAG2-null*^*A*^ (Figure 1G).

To determine if *STAG2* mutation affects *RUNX1* and *ERG* expression during megakaryocyte differentiation, we stimulated cells with PMA and used quantitative PCR to measure changes over 72 hours. Parental K562 cells showed gradual induction of *RUNX1-P1* and *ERG* transcription during stimulation (Supplementary Figure 12 and Figure 1H). In contrast, *STAG2*-*null* cells showed a precocious spike of *RUNX1* transcription 6-12 hours post-stimulation from its proximal P2 promoter (Figure 1I and Supplementary Figure 12). A similar precocious spike was observed in transcription of *ERG* (Figure 1H). By 48 hours post-stimulation, *RUNX1* and *ERG* transcription had returned to baseline in *STAG2*-*null* cells. These results imply that increased chromatin accessibility at *RUNX1* and *ERG* in *STAG2-null* cells leads to unrestrained transcription in response to differentiation stimuli. K562 parental cells upregulated *GATA1* and downregulated *KLF1* by 48 hours post-stimulation (Supplementary Figure 13), consistent with megakaryocyte differentiation. While *STAG2*-*null* cells successfully downregulated *KLF1*, they were not able to upregulate *GATA1.*

BRD4 is a bromodomain-containing protein that associates with active enhancers (Bhagwat et al., 2016). Notably, BRD4 binds at the *RUNX1-P2* and *ERG* +85 enhancer (Figure 1G). JQ1 is a BET inhibitor protein that reduces BRD4 binding and dampens SE-driven transcription. BRD4 can be removed from *RUNX1* and *ERG* by the BET inhibitor, JQ1 (Figure 1G, data from (Liu et al., 2017)). We treated *STAG2*-*null* cells with JQ1 together with PMA, and measured expression spikes in *RUNX1-P2* and *ERG*. JQ1 reduced *RUNX1-P2* and *ERG* expression in parental cells and strikingly, dampened the PMA induced transcription spikes seen in *STAG2-null* cells (Figure 1H-I and Supplementary Figure 12). *RUNX1-P1* transcription was completely blocked by JQ1 in both WT and *STAG2-null* cells (Supplementary Figure 12).

*STAG2-null* cells have reduced expression of the differentiation marker CD15 and elevated levels of the stem cell-associated marker, KIT (CD117), which is only lowly expressed in K562 cells (Figure 1J and Supplementary Figure 14A). Following 24 hours of treatment with JQ1, cell surface protein levels of KIT reduced by 2-fold in both *STAG2-null* clones while mRNA was reduced dramatically following 6 hours of treatment (Figure 1J and Supplementary Figure 14A-B). However, JQ1 treatment did not increase CD15 in *STAG2-null* cells (Figure 1J and Supplementary Figure 14A) implying that differentiation is not rescued. Collectively, the data indicate that BET inhibition can limit precocious *RUNX1/ERG* transcription and reduce leukemic stem cell-associated KIT expression in STAG2 mutant cells.

Overall our results suggest that cohesin-STAG2 depletion de-constrains the chromatin around *RUNX1* and *ERG*, which causes aberrant enhancer-amplified transcription in response to differentiation signals. We show that enhancer suppression using BET inhibitor, JQ1 prevents aberrant *RUNX1* and *ERG* signal-induced transcription in STAG2 mutant cells and reduces leukemic stem cell characteristics of STAG2 mutants.

## Acknowledgements and Author contributions

We would like to thank Catherine Young and Michelle Wilson from the Otago Flow cytometry facility (NZ) and Silke Newman for assistance and advice on flow cytometry. This work was supported by Health Research Council of NZ award 15/229 to J.A.H, and a Cancer Research Trust of NZ award to J.A and J.A.H. J.A. and J.A.H. designed research; J.A., T.T., U.K., R.D. and I.M.M. performed experiments; J.A., G.G. and J.A.H. analyzed data; J.A., G.G. and J.A.H. wrote the paper

