## Supplementary Material for "BET inhibition prevents aberrant RUNX1 and ERG transcription in STAG2 mutant leukaemia cells"

### Supplementary Methods

#### Cell culture

K562 cells were acquired from ATCC and maintained in Isocove's Modified Dulbecco's Media (IMDM) (Life Technologies) containing 10% fetal bovine serum. Adherent *STAG2-null* lines were detached for subculture using trypsin-EDTA (0.005% final concentration, Life Technologies). Differentiation to megakaryocytes was induced using 30 nM of phorbol 12-myristate 13-acetate (Kuvardina et al., 2015) (PMA, Sigma). BET inhibitor JQ1 (catalogue #S7110; TOCRIS) was used at 500 nM (Bhagwat et al., 2016). Images of cells in culture were taken using x20 objective on an Olympus THP4.

#### CRISPR-CAS9 editing

CRISPR-CAS9 editing was used to create STAG2 R614* mutation in K562 cells. pSpCas9(BB)-2A-GFP (PX458) was sourced from Addgene (plasmid #48138) (Ran et al., 2013). The sgRNA sequence (5'-ATTCCGGATCTGTCGCAATA-3') was designed using the http://crispr.mit.edu/ website and cloned into PX458 as previously described (Ran et al., 2013). A silent mutation (C>A) was introduced into the PAM site of the single stranded oligodeoxynucleotide (ssODN) donor repair template (IDT) template to prevent further cutting by Cas9. 4 million K562 cells were transfected with 20 µg of plasmid and 50 µl of 10 µM ssODN using a Neon transfection system (Life Technologies). Single GFP-positive cells were isolated using a FACSAriaII (Becton Dickinson). Clones were screened for the point mutation by restriction digest and Sanger sequencing. ssODN repair template sequence: ATATATGCCTTAGTTTTGATGAAAAATATTATTATTTTGCAGCATTTGGATGCATTATTGTGACAGATCCGGAATATTGTAGAGAAGCACACAGATACAGATGTTTTGGAAGCATGTTCTAAAACTTACCATGCACTCTGTAATGAAGAGTTCACAATCT; Green highlight: silent mutation (C>A) introduced into the PAM site; Yellow highlight: R614* (C>T) change.

#### RNA sequencing and analyses

Total RNA was extracted using NucleoSpin RNA kit (MACHEREY-NAGEL). Libraries from three biological replicates of each cell type were prepared using NEBNext® Ultra™ RNA Library Prep Kit (Illumina®) and sequenced on HiSeq X (150 bp paired end reads, 20 million reads per sample) by Annoroad Gene Technology Ltd. (Beijing, China), contracted through Custom Science (NZ). RNA sequencing reads were aligned to the human genome

GRCh37 (hg19) using HISAT2 version 2.0.5 with gene annotation from Ensembl version 75. Read counts were retrieved by exon and summarized by gene using featureCount version v1.5.3. Differentially expressed genes in the *STAG2-null* lines versus parental (WT) were identified using DESeq2 (Love et al., 2014). P-values were adjusted for multi test using Independent Hypothesis Weighting (Ignatiadis et al., 2016). Euclidean distance matrix correlation was computed in R, based on clustering of biological replicates for each cell type. GSEA software (Subramanian et al., 2005) (settings; enrichment statistics: weighted; max size: 5000) was used to carry out enrichment analyses on ranked gene lists based on Wald statistics from DESeq2 on MgSigDB gene sets. The ranked lists contained genes significant at adjusted p-values $\leq 0.05$ from the DESeq2 analyses.

**ATAC sequencing and analyses**

50,000 cells were transposed as described previously (Buenrostro et al., 2015; Litzenburger et al., 2017). Libraries from three biological replicates of parental K562 (WT) and *STAG2-null*$^{A}$ were prepared using Illumina® Nextera DNA library preparation (FC-121-1030) and indexing (FC-121-1011) kits, and sequenced at 100 bp single end reads on HiSeq 2500 at the Otago Genomics Facility (New Zealand). ATAC-sequencing reads were aligned to the human genome GRCh37 (hg19) using bowtie2. Peaks were called using Macs2 without model and shift size to accommodate transposase integration (--nomodel --shift -37 --extsize 73). Peak summits were extended ±250 bp. Peaks from all biological replicates were merged and normalized. Only peaks overlapping in all 3 replicates were considered. Biological replicates correlation matrix was computed using Euclidean distance.

**ATAC sequencing differential analyses**

Differential peaks were identified using edgeR (McCarthy et al., 2009). Briefly, peak count data were retrieved using featureCounts (Smyth et al., 2013). Differential events were estimated using Generalized Linear Models. After likelihood test ratio, p-values were adjusted for multiple testing using Benjamini-Hochberg FDR <0.05.

**ATAC sequencing peak annotation**

Peaks were annotated to genomic sites and genes using HOMER with Hg19 annotation (Heinz et al., 2010). Enrichment of differential ATAC peaks at super enhancers (dbsuper database; (Khan and Zhang, 2016)) was determined using the 'intersectBED' function from BED tools utilities (Quinlan and Hall, 2010).

**ATAC sequencing motif analyses**

Enrichment of transcription factor binding motifs were analysed using HOMER for known motifs (Heinz et al., 2010).

**Array CGH**

Array comparative genomic hybridization was performed using the Agilent SurePrint G3 ISCA CGH+SNP 4x180k(hg19) array platform at the Wellington Genetic Services Facility (New Zealand). The parental K562 was used as the reference for array CGH.

**Quantitative PCR (RT-qPCR).**

cDNA from total RNA was made using Qscript™ cDNA SuperMix (Quantbio). RT-qPCR was performed on a LightCycler® 480 II (Roche Life Science) using SYBR® Premix Ex Taq™ (Takara). Expression values relative to reference genes cyclophilin and glyceraldehyde 3-phosphate dehydrogenase (GAPDH) were derived using qBase Plus (Biogazelle). Primer sequences are provided in Supplementary Table 1.

**Immunoblotting**

Immunoblotting was carried out as described previously (Antony et al., 2015; Dasgupta et al., 2016). Antibodies were anti-STAG2 (ab4463, Abcam, 1:1000) and anti-γ-Tubulin (T5326, Sigma, 1:5000), detected using IRDye 800CW Donkey anti-Goat IgG and IRDye 680CW Goat anti-mouse IgG (LICOR) respectively. LI-COR Odyssey® and LI-COR Image Studio software was used to image and quantify blots.

**Flow cytometry**

Cells treated with 500 nM JQ1 or DMSO were stained with PE anti-CD117 (KIT) (340529, BD Biosciences) and APC anti-CD15 (551376, BD Biosciences), flow cytometry was carried out on the Navios EX flow cytometer at Southern Community Laboratories (New Zealand) and analysed using Kaluza Software (Beckman Coulter). Cell cycle progression was analysed following double thymidine block and propidium iodide as previously described (Antony et al., 2015; Dasgupta et al., 2016) and analysed using Flow-JO software.

**Statistical analyses**

Unless otherwise stated all statistical analyses were carried out using Prism version 7 software (GraphPad).

### Data availability and datasets used

All data generated in this study have been deposited in NCBI's GEO database under the SuperSeries GSE131449 (RNA sequencing – GSE131448; ATAC sequencing – GSE131447). Datasets for BRD4 (GSE88817), FOS (wgEncodeEH000619), FOSL1/FRA1 (wgEncodeEH001637), JUN (wgEncodeEH000620) and JUND (wgEncodeEH002164) binding in K562 cells were used. Super enhancer bed files for K562, human CD14+ monocytes and human CD34+ primary cord blood cells (RO01536) were extracted from the dBsuper database  (Khan and Zhang, 2016).

### Supplementary Table 1: RT-qPCR primer sequences

| **Gene** | **Forward (5'-3')** | **Reverse (5'-3')** |
|---|---|---|
| *CYCLOPHILIN* | ACGGCGAGCCCTTGG | TTTCTGCTGTCTTTGGGACCT |
| *GAPDH* | TGCACCACCAACTGCTTAGC | GGCATGGACTGTGGTCATGAG |
| *STAG2* | ACGGAAAGTGGTTGAGGG | GTGGAGGTGAGTTGTGGTGT |
| *RUNX1_common* | AGACCCTGCCCATCGCTTTC | GGTTCTTCATGGCTGCGGTA |
| *RUNX1_P1* | TTTTCAGGAGGAAGCGATGG | TGGCATCGTGGACGTCTCTA |
| *RUNX1_P2* | ATAAAGGCCCCTGAACGTG | GTTAGGACCCTGCAAACAGC |
| *ERG* | CGCAGAGTTATCGTGCCAGCAGAT | CCATATTCTTTCACCGCCCACTCC |
| *GATA1* | TTGCCACATCCCCAAGGCGG | GGGGGAGGGGCTCTGAGGTC |
| *KLF1* | TGACTTCCTCAAGTGGTGGC | GGTGAGGAGGAGATCCAGGT |
| *KIT* | CAGGAAGATCATGCAGAAGCTG | TATCGCTGCAGGAAGACTCC |

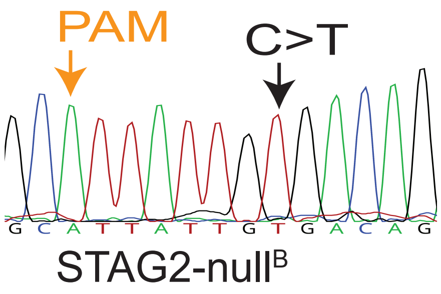


**Supplementary Figure 1**. Sanger sequencing plot of CRISPR-Cas9 edited K562 line homozygous (*STAG2-null*$^{B}$) for STAG2 R614*. The silent mutation introduced at PAM site is also indicated.

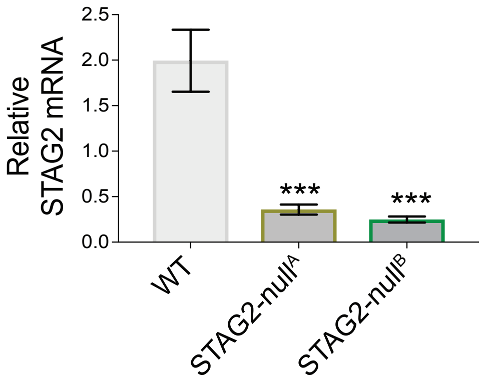


**Supplementary Figure 2**. Quantitative RT-PCR of *STAG2* mRNA in the parental (WT), and *STAG2-null* clones. Graph depicts *STAG2* mRNA from 3 biological replicates relative to 2 reference genes (***p<0.001; one-way Anova).

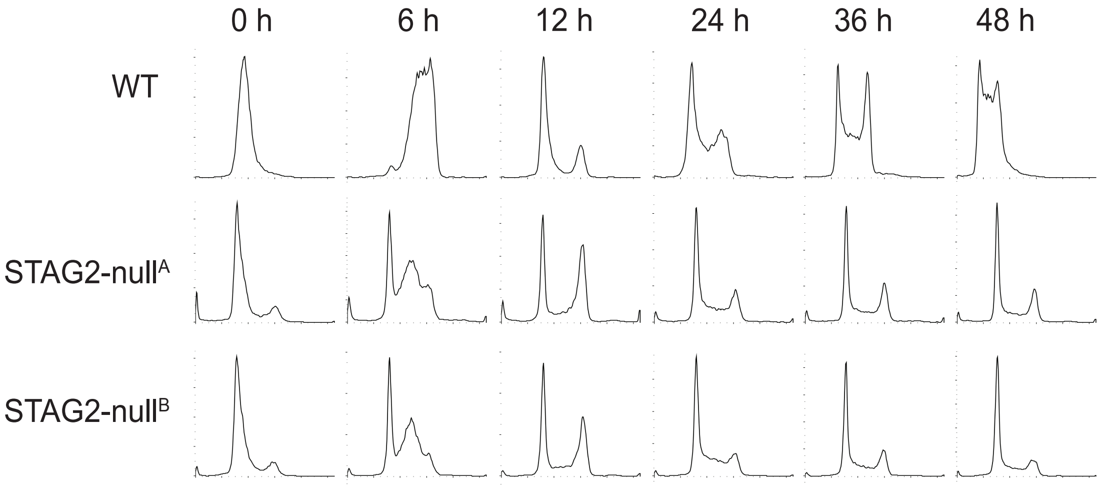


**Supplementary Figure 3**. Cell cycle analyses in parental (WT) and *STAG2-null* cells. Cells were synchronized by double thymidine block and then released and stained with propidium iodide at the indicated time points (Antony et al., 2015; Dasgupta et al., 2016). Cell cycle distributions were analyzed by flow cytometry on Beckman Coulter Gallios flow cytometer and using Flow-JO software. Histograms show propidium iodide staining intensity on the X-axis and cell counts on the Y-axis. *STAG2-null* cells show delay in cell cycle progression.

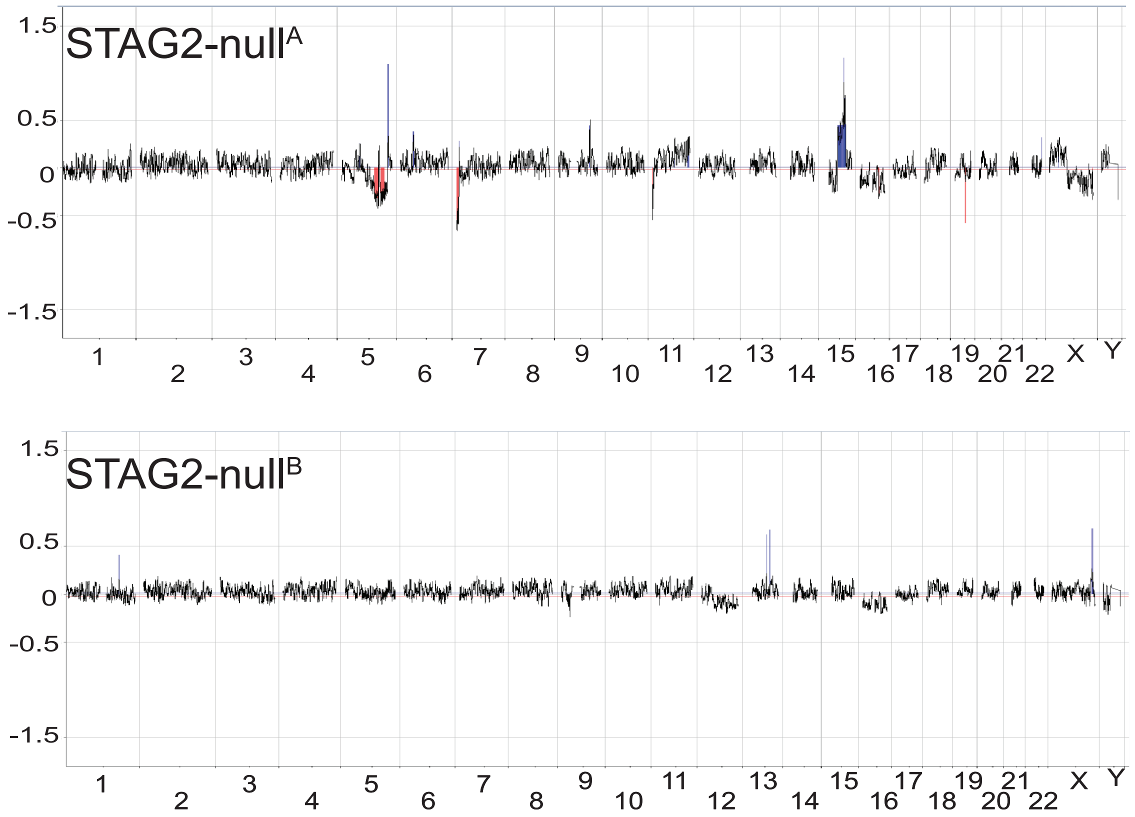


**Supplementary Figure 4.** Array CGH on *STAG2-null* cell lines. Multiple regions of mosaic gains and losses were detected in both *STAG2-null* lines. Relevant to this study, note that there are no gains or losses of Chromosome 21. Chromosome numbers are indicated on the X-axis and the Y-axis shows the log2 scores of gain or loss in *STAG2-null* lines compared to parental K562. Data was analyzed and visualized using the Agilent CytoGenomics Software.

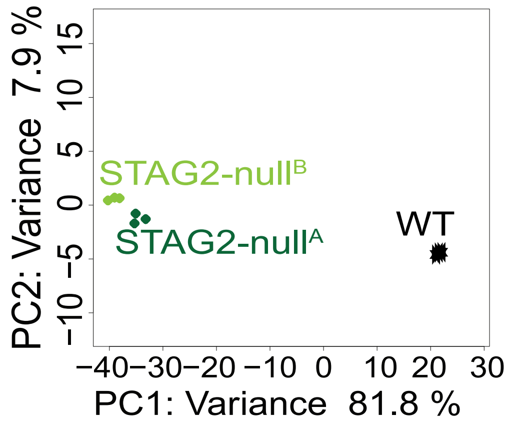


**Supplementary Figure 5.** Principal component analyses (PCA) of RNA sequencing outputs from parental (WT) and *STAG2-null* cells. Transcriptome profile of both *STAG2-null* lines are similar and cluster seperately from the parental K562 cells.

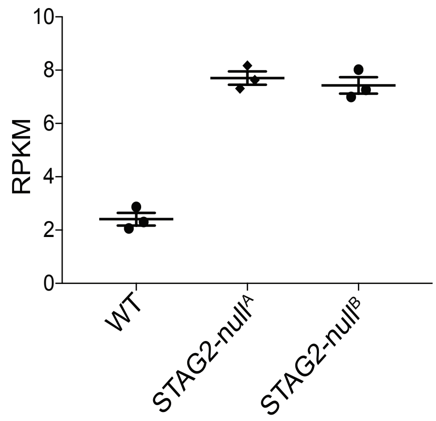


**Supplementary Figure 6**. Reads per kilobase million (RPKM) values of *STAG1* mRNA expression derived from RNA sequencing of parental (WT) and *STAG2-null* K562 cell lines. *STAG2-null* lines show significant ($p<0.0001$) 1.6-fold increase in *STAG1* mRNA levels compared to parental cells.

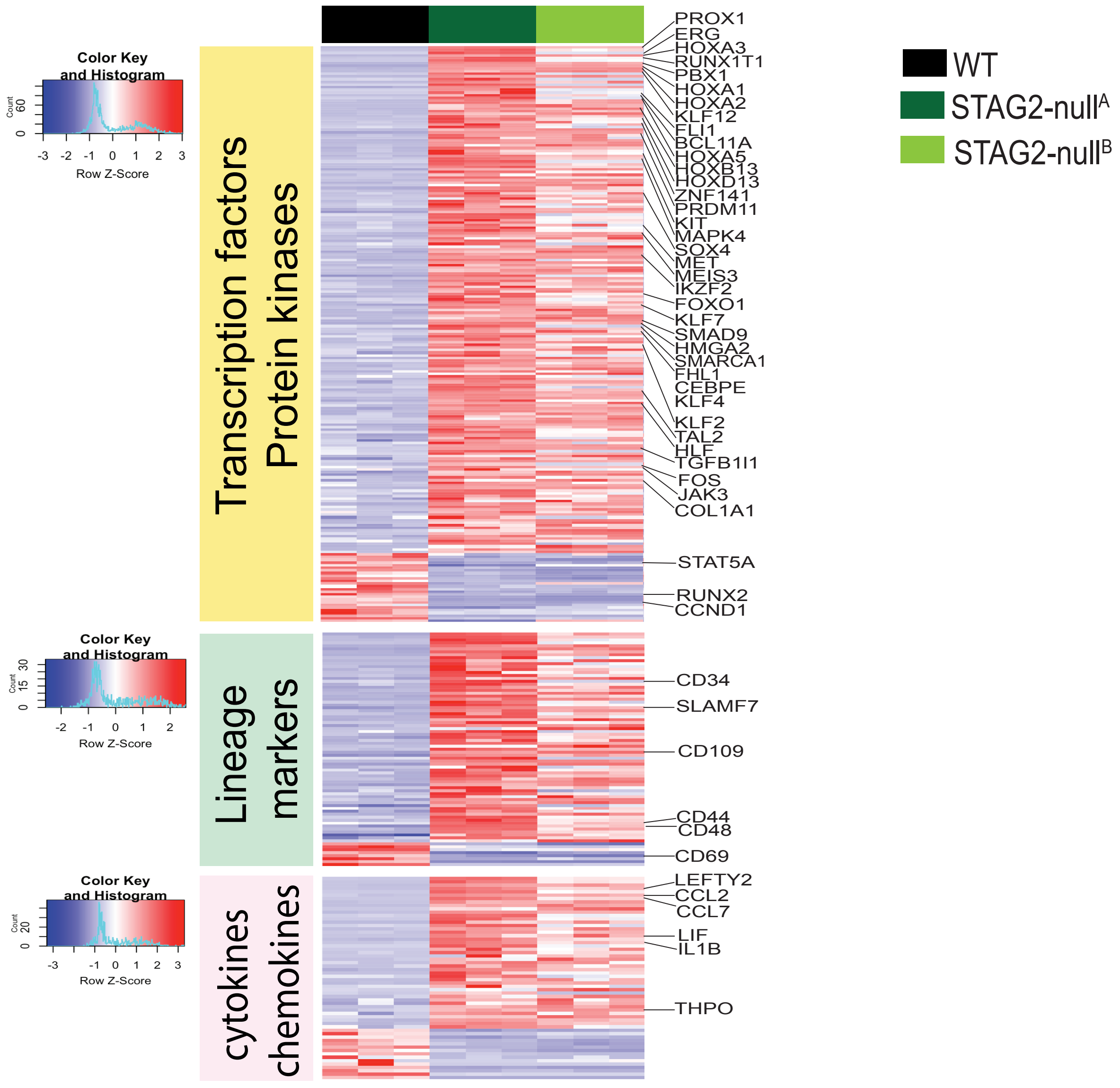


**Supplementary Figure 7.** Heat map showing log2 RPKM values derived from RNA sequencing. Genes shown are from the molecular signature database (MSigDB) identification of families of transcription factors, kinases, lineage markers, chemokines and cytokines that are significantly differentially regulated (adjusted p value $\leq 0.05$; log2 fold change $\geq 2$) in *STAG2-null^A* compared to parental (WT).

### GSEA STAG2-null cells

| Gene set | STAG2-null[A] | STAG2-null[B] |
| --- | --- | --- |
| Diaz chronic myelogenous leukemia_down (genes downregulated by BCR-ABL in CD34+ cells) | 2.15 (0.002) | 2.06 (0.019) |
| Diaz chronic myelogenous leukemia_up (genes upregulated by BCR-ABL in CD34+ cells) | -2.45 (0.0002) | -2.33 (0.043) |
| Naba core matrisome | 2.47 (0) | 2.41 (0) |
| Naba ECM glycoproteins | 2.38 (0.00014) | 2.43 (0) |
| Hallmark epithelial mesenchymal transition | 2.94 (0) | 2.15 (0.00024) |
| Gal leukemic stem cell_up | 1.8 (0.0028) | 1.65 (0.092) |
| Jaatinen hematopoietic stem cell_up | 1.8 (0.0184) | 1.45 (1.98) |
| Eppert HSC_R (genes upregulated in HSC compared to differentiated cells) | 2.01 (0.007) | 1.46 (0.194) |

**Supplementary Figure 8.** Gene set enrichment analyses (GSEA) of significant differentially expressed genes in *STAG2-null* cells using gene sets from the molecular signature database (MSigDB). The normalized enrichment score (NES) for each group is shown (FDR q value). Positive NES indicates enrichment for upregulation and negative NES indicates enrichment for downregulation of genes for the analyzed gene set.

Diaz Chronic myelogenous leukemia_down: Genes normally dowregulated by BCR-ABL are upreuglated in *STAG2-null* cells.

Diaz Chronic myelogenous leukemia_up: Genes normally upregulated by BCR-ABL are down-regulated in *STAG2-null* cells.

Extracellular matrix genes (Naba core matrisome, Naba ECM glycoproteins, Hallmark epithelial mesenchymal transition) are upregualted in *STAG2-null* cells.

Stem cell associated genes (Gal leukemic stem cell_up, Jaatinen hematopietic stem cell_up and Eppert HSC_R) are upregualted in *STAG2-null* cells.

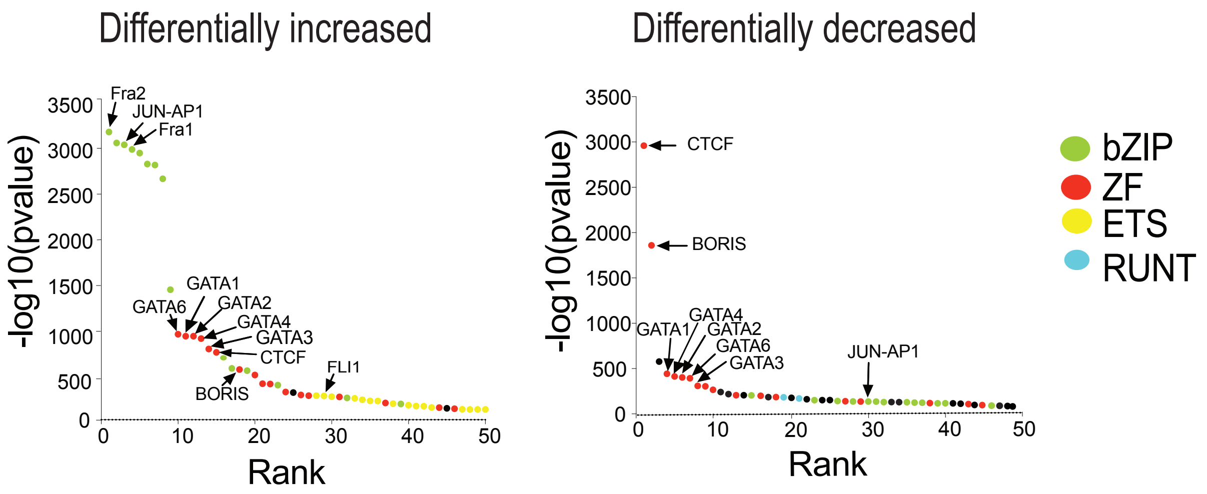


**Supplementary Figure 9.** Motif enrichment analyses using HOMER at differentially accessible ATAC sites in *STAG2-null*[A]. Graphs show log10 p-value on the Y axis and ranking of the motif based on significance (highest to lowest) on the X axis. The bZIP, ZF, ETS and RUNT families are indicated in green, red, yellow and blue respectively.

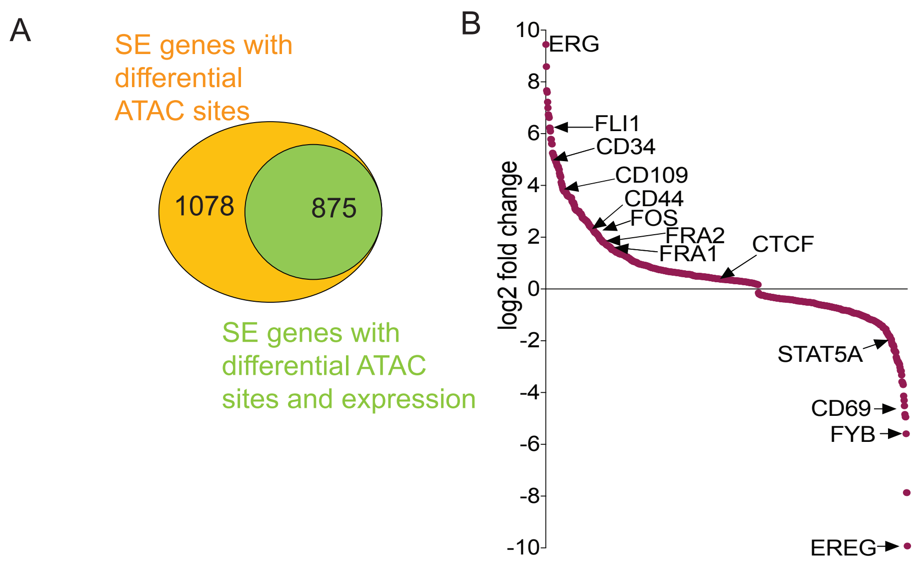


**Supplementary Figure 10.** (A) Venn diagram depicting the overlap of genes in proximity to differentially accessible SEs in *STAG2-null[A]* that also show differential expression. (B) Differentially expressed SE genes that have differential chromatin accessibility in *STAG2-null[A]* ($p \leq 0.05$). Graph shows ranking based on log2 fold change from RNA sequencing analyses.

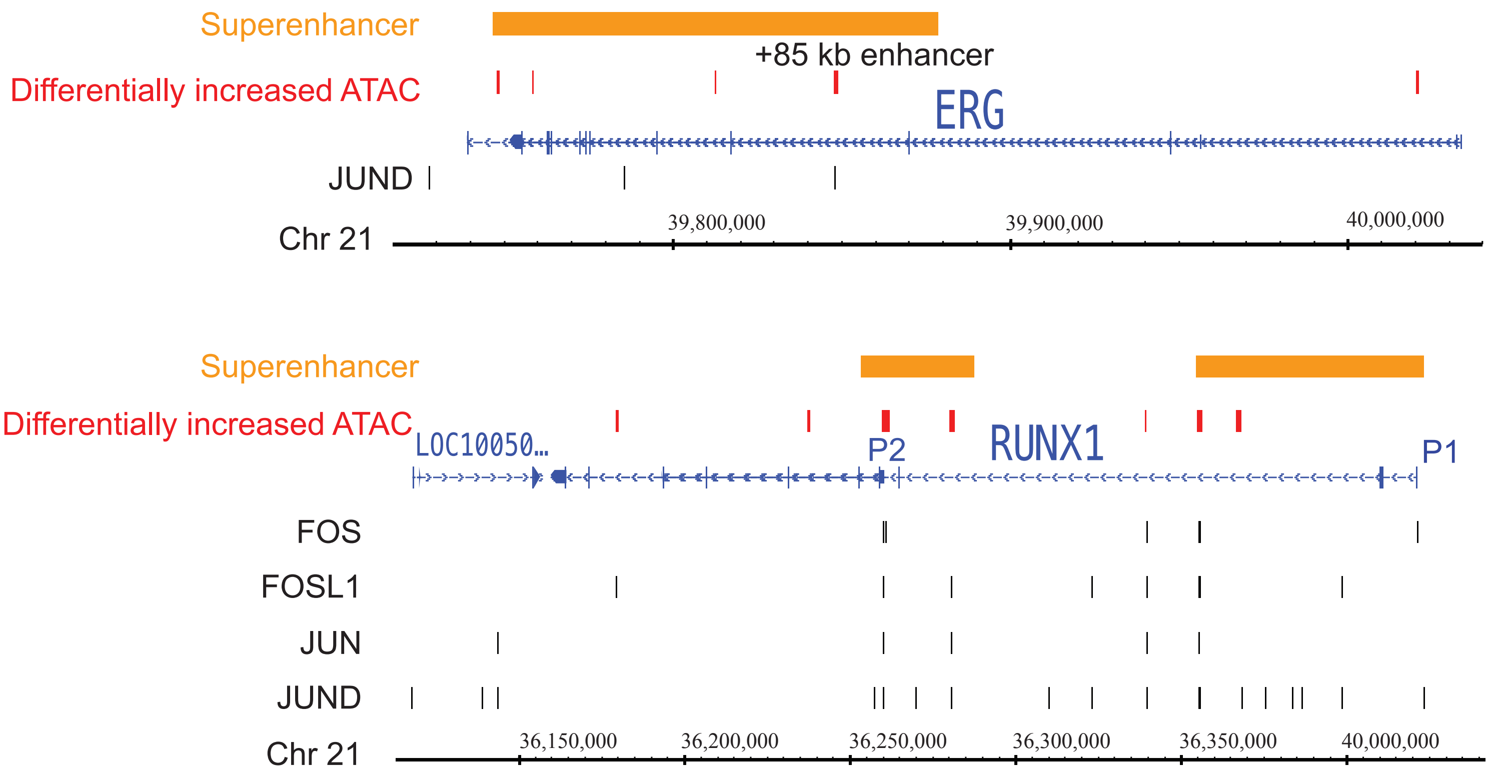


**Supplementary Figure 11**. *ERG* and *RUNX1* genomic regions. Shown are tracks of superenhancer sites in CD34+ primary cord blood cells (RO01536; derived from dB superenhancer database; Khan and Zhang, 2016). Differentially increased ATAC sites identifed in *STAG2-null[A]* cells are shown in red. Binding sites of AP-1 factors FOS, FOSL1 or FRA1, JUN and JUND in K562 are also indicated. Binding sites for AP-1 factors are derived from ENCODE ChIP sequencing datasets.
FOS (wgEncodeEH000619), FOSL1/FRA1 (wgEncodeEH001637), JUN (wgEncodeEH000620) and JUND (wgEncodeEH002164) .

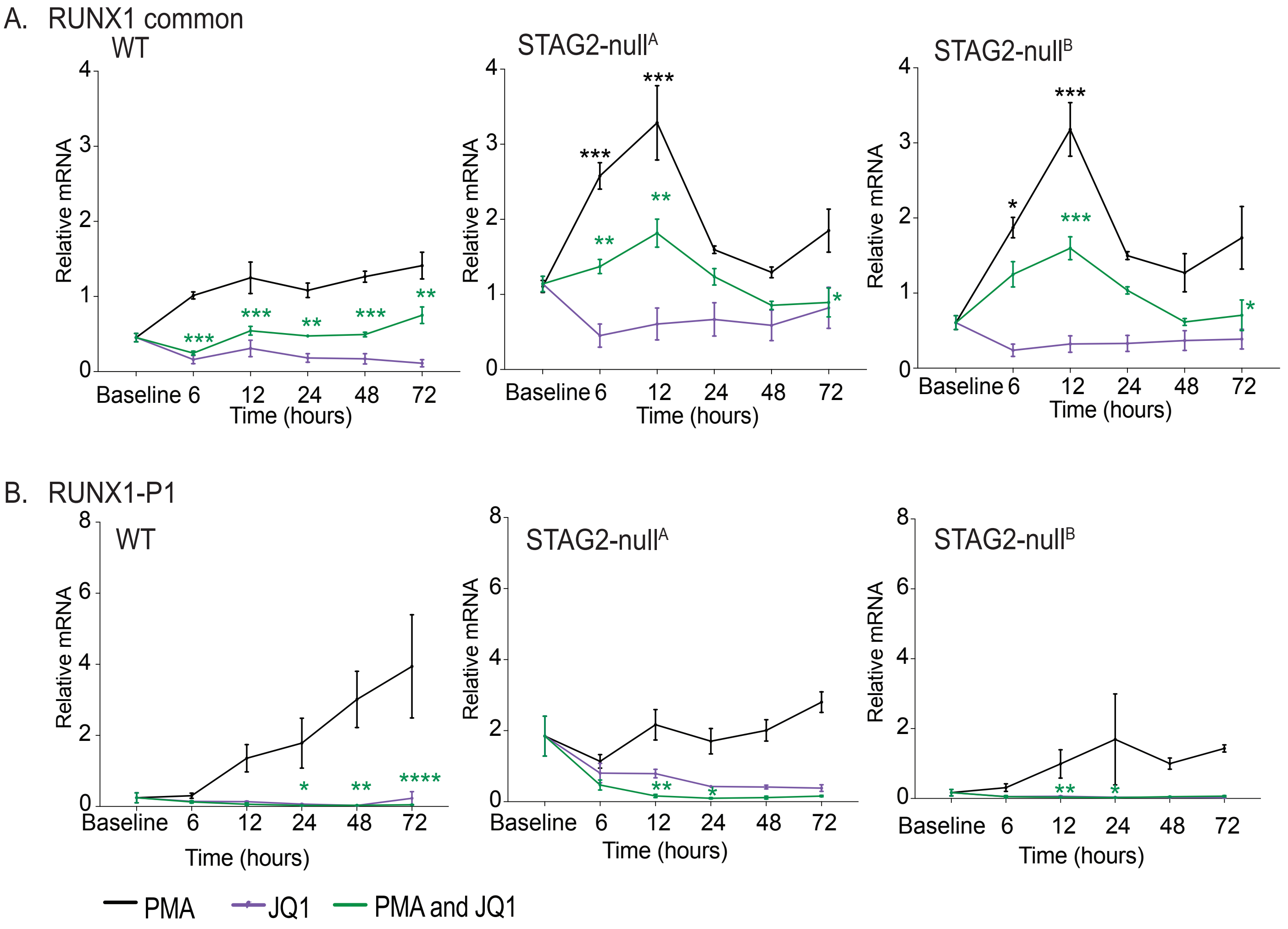


**Supplementary Figure 12.** RT-qPCR analyses using primers to detect (A) common *RUNX1* transcript and (B) *RUNX1-P1* transcript at the indicated time points after treatment with PMA, JQ1 or a combination of PMA and JQ1. Black asterisks denote significant difference between parental (WT) and *STAG2-null* lines at the specific time point following PMA only treatment. Green asterisks denote significant difference between PMA only and combination of PMA and JQ1 within each cell type. Significance was determined by two-way Anova ( $^*p< 0.05$, $^{**}p<0.01$, $^{***}p<0.001$, $^{****}p<0.0001$).

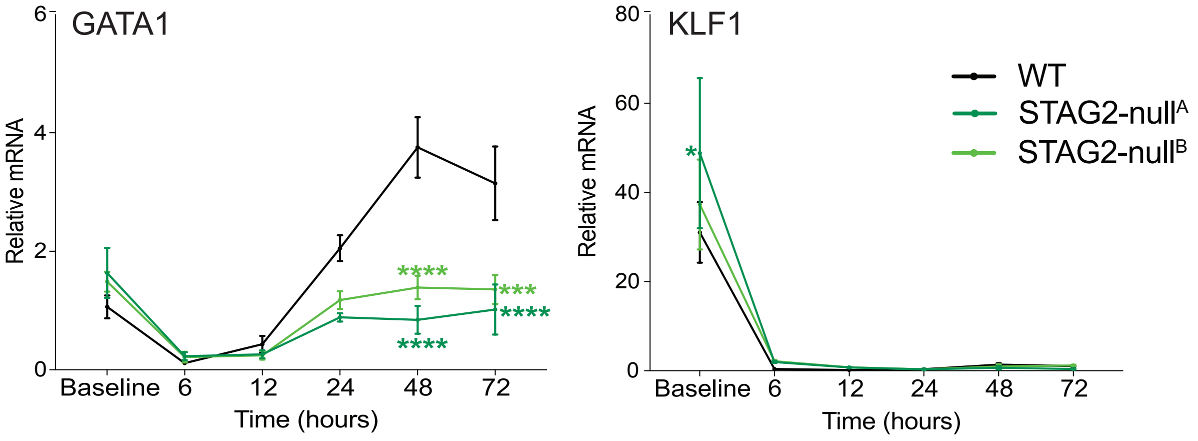


**Supplementary Figure 13.** RT-qPCR analyses using primers to detect *GATA1* and *KLF1* transcripts at the indicated time points after treatment with PMA. Asterisks denote significant change from parental cells (WT). Significance was determined by two-way Anova ( $^*p<0.05$, $^{***}p<0.001$, $^{****}p<0.0001$).

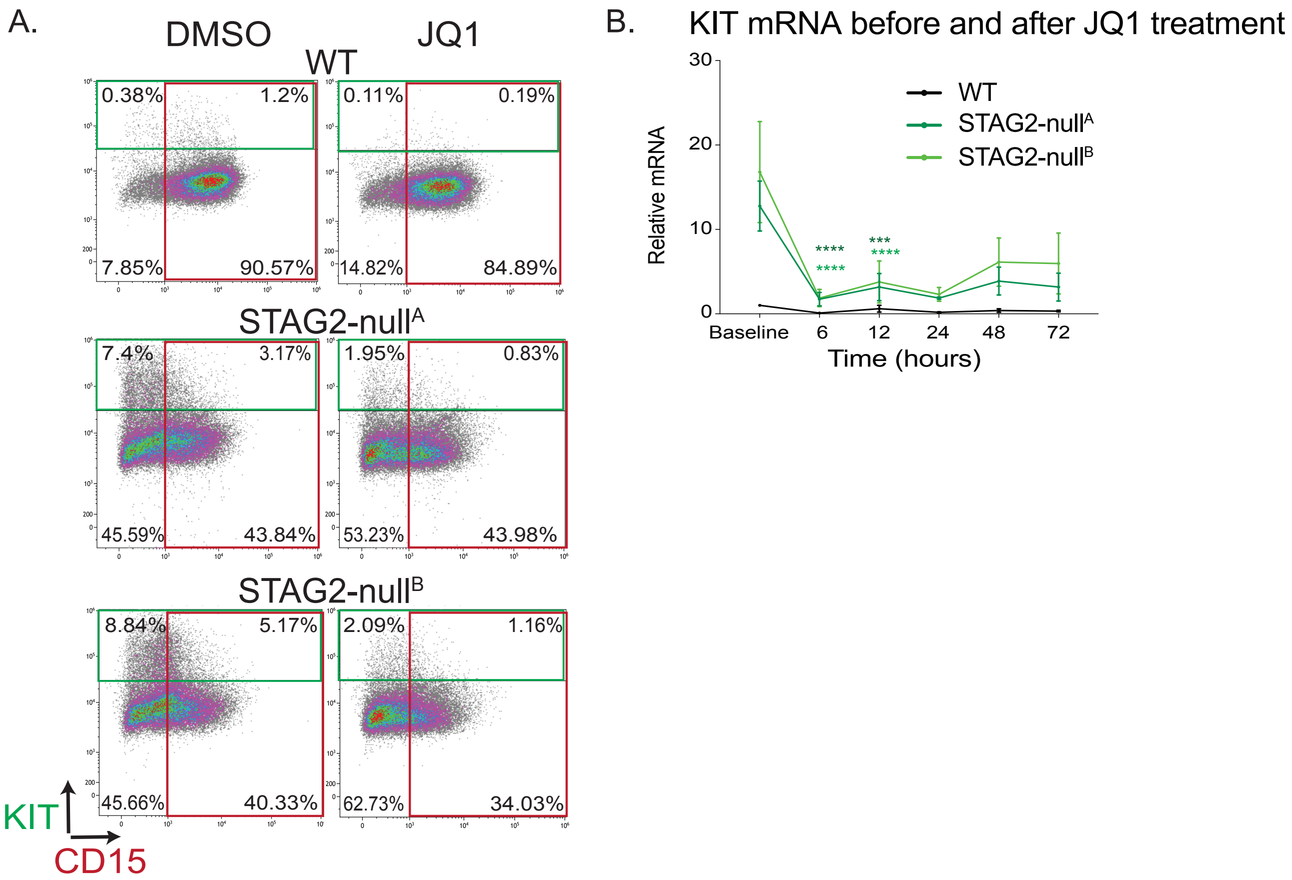


**Supplementary Figure 14.** (A) Representative plots of flow cytometry analyses of KIT and CD15 markers in *STAG2-null* and parental (WT) K562 cells following 24 hours of DMSO (control) or JQ1 treatment. The fluorescence intensity of KIT is on the Y axis and CD15 on the X axis. (B) RT-qPCR analyses of *KIT* transcript at various time points following JQ1 treatment. The graph represents the average *KIT* transcript levels from 3 biological replicates normalized to 2 reference genes. Asterisks denote significant change from baseline levels within each cell type. Significance was determined by two-way Anova ( ***p<0.001, ****p<0.0001).
